# SARS-CoV-2 assays to detect functional antibody responses that block ACE2 recognition in vaccinated animals and infected patients

**DOI:** 10.1101/2020.06.17.158527

**Authors:** Susanne N. Walker, Neethu Chokkalingam, Emma L. Reuschel, Mansi Purwar, Ziyang Xu, Ebony N. Gary, Kevin Y. Kim, Katherine Schultheis, Jewell Walters, Stephanie Ramos, Trevor R.F. Smith, Kate E. Broderick, Pablo Tebas, Ami Patel, David B. Weiner, Daniel W. Kulp

**Affiliations:** Vaccine and Immunotherapy Center, Wistar Institute, Philadelphia, PA, 19104, USA; Inovio Pharmaceuticals, Plymouth Meeting, PA, 19462, USA; Department of Medicine; University of Pennsylvania Perelman School of Medicine, Philadelphia, PA, 19104, United States; Department of Microbiology, Perelman School of Medicine, University of Pennsylvania, Philadelphia, PA 19104, United States

**Keywords:** SARS-CoV-2, COVID-19, ACE2 blocking assay, Serological tests, Functional antibodies, SARS-CoV-2 vaccine, SARS-CoV-2 immunity

## Abstract

SARS-CoV-2 (<u>S</u>evere <u>A</u>cute <u>R</u>espiratory <u>S</u>yndrome <u>Co</u>rona<u>v</u>irus <u>2</u>) has caused a global pandemic of COVID-19 resulting in cases of mild to severe respiratory distress and significant mortality. The global outbreak of this novel coronavirus has now infected >8 million people worldwide with >2 million cases in the US (June 17^th^, 2020). There is an urgent need for vaccines and therapeutics to combat the spread of this coronavirus. Similarly, the development of diagnostic and research tools to determine infection and vaccine efficacy are critically needed. Molecular assays have been developed to determine viral genetic material present in patients. Serological assays have been developed to determine humoral responses to the spike protein or receptor binding domain (RBD). Detection of functional antibodies can be accomplished through neutralization of live SARS-CoV2 virus, but requires significant expertise, an infectible stable cell line, a specialized BioSafety Level 3 (BSL-3) facility. As large numbers of people return from quarantine, it is critical to have rapid diagnostics that can be widely adopted and employed to assess functional antibody levels in the returning workforce. This type of surrogate neutralization diagnostic can also be used to assess humoral immune responses induced in patients from the large number of vaccine and immunotherapy trials currently on-going. Here we describe a rapid serological diagnostic assay for determining antibody receptor blocking and demonstrate the broad utility of the assay by measuring the antibody functionality of sera from small animals and non-human primates immunized with an experimental SARS-CoV-2 vaccine and using sera from infected patients.

## Background

The city of Wuhan China became the epicenter for a global pandemic in December of 2019 when the first cases of respiratory illnesses were reported and identified as being caused by a novel betacoronavirus now referred to as SARS-CoV-2. The novel SARS-CoV-2 virus is closely related to the SARS-CoV-1 virus and has a high human-to-human transmission rate(1). As of May 22^nd^ 2020, over 5,128,492 people are reported positive for viral infection of SARS-CoV-2 resulting in 333,489 fatalities worldwide(2). SARS-CoV-2 modes of transmission include shedding of the virions in airborne droplets and close contact. Once infected, people develop flu-like symptoms and severe cases can lead to acute respiratory distress syndrome (ARDS), acute respiratory failure, severe pneumonia, pulmonary edema, multiple organ failure and death. Quarantines around the globe have helped curb spread of the virus and as people are eager to return to life as normal, development of methods and assays to aid in the detection of SARS-CoV-2 and effective measurements are of utmost importance.

Angiotensin-converting enzyme 2 (ACE2) is highly expressed on lung epithelial cells and is the viral receptor for SARS-CoV-1(3). Recently, ACE2 has also been identified as a receptor utilized by SARS-CoV-2(4). A 193 amino acid region on the spike protein of SARS-CoV-1 and SARS-CoV-2 termed the receptor binding domain (RBD) was found to interact with the ACE2 receptor and primarily mediates cell entry(5–10). Cryo-electron microscopy was used to determine a high resolution structure of the prefusion spike protein and help define the ACE2 interaction site(11, 12). In the SARS-CoV-1 outbreak of 2003, it was reported that high titers of protecting IgG were found in the convalescent serum of patients recovering from infection(13). Early studies show treating COVID-19 patients presenting with severe ARDS with convalescent plasma therapy (CPT) may be beneficial(14, 15). In order to determine if a recovered patient could be re-infected or if their plasma may be useful as a treatment to others, assessment of functional antibody response is highly valuable. While tests for SARS-CoV-2 seropositivity are being deployed, they do not necessarily correlate with neutralizing immunity(16). As there are limited options for rapid diagnostic of functional antibody responses, fast and simple functional assays may prove to be a critical assessment tool to discriminate between potential CPT donors.

In parallel with CPT, major academic, industry and government entities are pushing for therapeutics and vaccines against SARS-CoV-2. Given this urgent need, there are numerous clinical vaccine trials underway for SARS-CoV-2 in parallel(17). Through pre-clinical testing, vaccine-induced functional antibody responses are being used to discriminate among potential vaccine candidates. In clinical testing, the humoral responses will need to be analyzed for functionality to assess immunogenicity of each vaccine. Therefore, rapid assays for detecting functional IgGs in human serum for SARS-CoV-2 are essential to compare vaccine candidates and to understand induced immunity.

Early into this pandemic, diagnostic tools for detecting viral genetic material were developed and utilized in a real-time reverse transcription PCR assay(18). Serologic assays for detecting anti-SARS-CoV-2 antibodies are now also available(19). However, single antigen ELISA assays can suffer from detection limits and numbers of false positives(20) and may not be associated solely with protective responses. Faster and straight forward approaches which detect specific interactions to reduce false positives are needed. Neutralization assays that can detect functional immunity have been developed with replicating virus, but require handling in specialized BSL-3 facilities, severely limiting the number of samples that can be processed. Pseudovirus neutralization assays run in BSL-2 facilities were quickly developed to detect the functional antibody response in sera(21). While this is a critical tool for determining protective antibody titers, it requires several days for a readout and are not standardized between laboratories. The pseudoviruses produced in these assays are not easily manufactured and take time to express, harvest, and titer. One such approach to help augment the methods listed above is an enzyme-linked immunosorbent assay (ELISA) employed in a competitive manner to determine levels of ACE2 receptor blocking antibodies in a sample. In addition, recent advances of portable and field-deployable surface plasmon resonance (SPR) devices(22) and widespread availability of SPR instruments in research laboratories make SPR an additional platform for measuring ACE2 receptor blocking. Here, we describe a competition ELISA assay and a SPR assay developed to rapidly detect ACE2 receptor blocking antibodies in IgGs and sera of vaccinated mice, guinea pigs, rabbits and non-human primates, as well as, human samples from SARS-CoV-2 patients.

## Results

### Receptor expression and assay development

To detect SARS-CoV-2 spike protein binding to ACE2, we designed a soluble variant of the membrane-bound human ACE2 receptor. The ectodomain of the receptor was fused to a human IgG1 Fc tag (ACE2-IgHu), for purification and secondary antibody recognition. This protein fusion was expressed in mammalian cells to ensure proper glycosylation and purified on a protein A column. To determine that ACE2-IgHu was dimeric and a homogeneous species in solution, we employed size exclusion chromatography tandem multi angle light scattering (SEC-MALS). The SEC trace shows a single peak (Fig. 1A) and the MALS data determined the estimated molecular weight to be 190 kDa for ACE2-IgHu, which is very close to the expected molecular weight of a dimer of ACE2-IgHu at 195 kDa (**Fig. 1B**). We next sought to confirm the functionality of ACE2-IgHu. Previous studies suggest SARS-CoV-2 binds to ACE2 with an affinity range of 4-34nM(6, 11). We determined that our ACE2-IgHu binds with similar affinity to the receptor binding domain (RBD) of SARS-CoV-2 spike (27.5nM) as assessed by SPR (**Fig. 1C**). Next, using Enzyme-Linked Immunosorbent Assays (ELISAs) we immobilized full-length SARS-CoV-2 spike protein (containing both the S1 and S2 subunits) and incubated a dilution series of ACE2-IgHu (**Fig. 1D**). The binding curves confirmed the high affinity interaction of the receptor for the spike protein. We further showed similar binding for two independent batches of ACE2, as well as a sample that was frozen and thawed (**Fig. S1A**). From the binding curve, we hypothesized that a reasonable concentration of ACE2 protein fusion needed to see competitive blocking while still binding >90% of immobilized spike protein in the absence of blocking would be around 0.1-1 ug/ml (**Fig. 1D, red arrow**).

**Figure 1.**
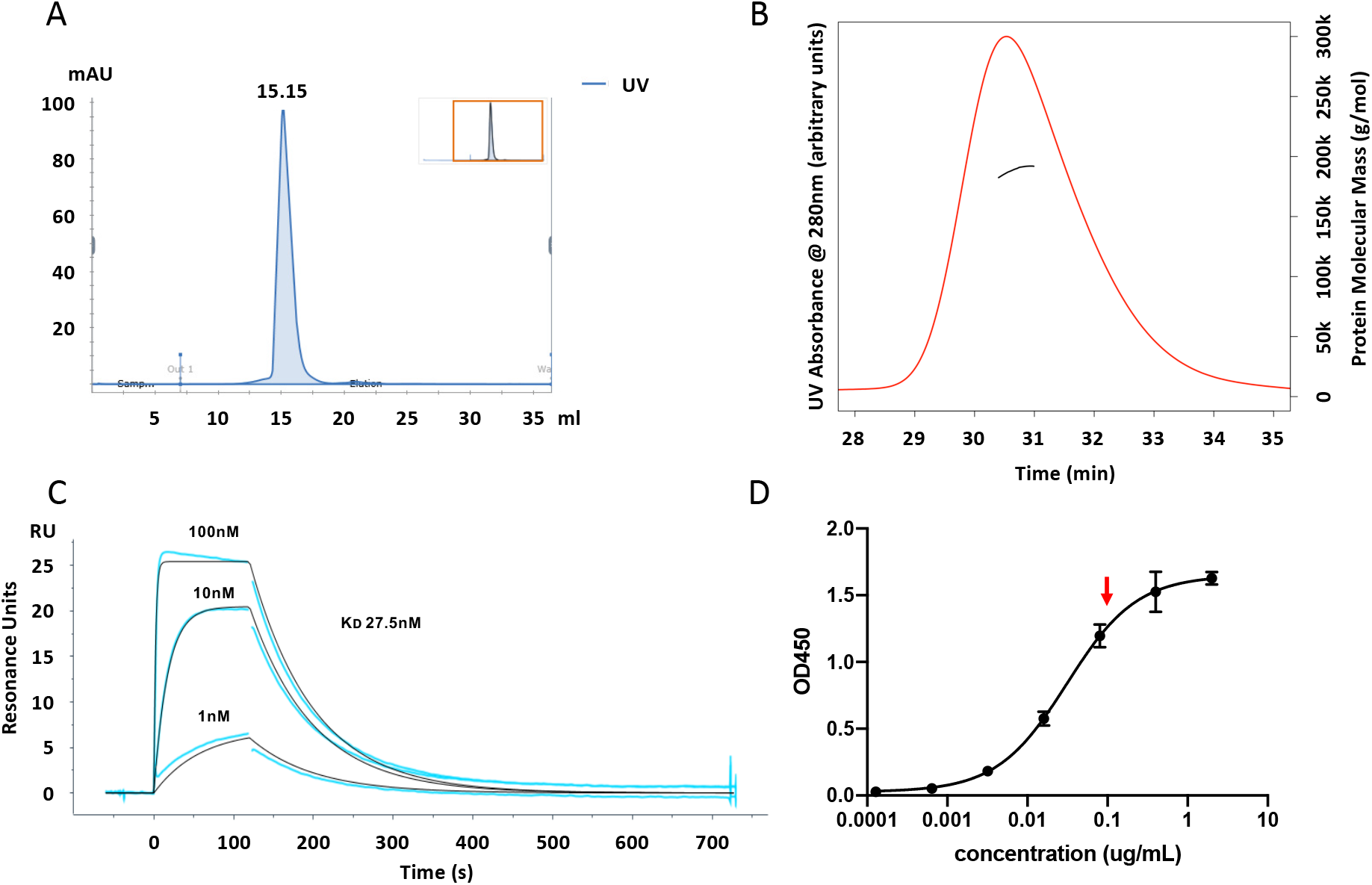
ACE2 receptor expression and affinity. **a**) UV trace from SEC of ACE2-IgHu expressed in mammalian cells and purified on a protein A column **b**) SEC-MALS determined molecular weight of ACE2-IgHu to be that of a dimeric complex (~190kDa) as shown by the line under the peak on the right y-axis **c**) Affinity of SARS CoV-2 receptor binding domain for immobilized ACE2-IgHu assessed by SPR (27nM) curves are concentrations of RBD X, Y, and Z **d**) Affinity of ACE2-IgHu for immobilized SARS CoV-2 full-length spike protein assessed by ELISA. Optimal concentration of ACE2-IgHu for competition assays (~EC_90_, red arrow) requires high signal without excess receptor present

To examine if we could construct a competition assay, we employed ACE2-IgMu (mouse Fc) to act as a competitor to ACE2-IgHu. To match our initial binding ELISA format, the competition ELISA assay similarly captured a His6X-tagged full-length spike protein by first immobilizing an anti-His polyclonal antibody. A dilution series of the competitor (ACE2-IgMu) was pre-mixed with a constant concentration of soluble receptor (ACE2-IgHu). A secondary anti-human antibody conjugated to horseradish peroxidase (HRP) determines the amount of ACE2-IgHu present through a TMB colorimetric readout (**Fig. 2A**). In order to formally determine the optimal concentration of soluble receptor (ACE2-IgHu), we performed the assay at four concentrations ranging from 0.01ug/ml to 10 ug/ml (**Fig. 2B**). The ACE2-IgHu concentration of 0.1 ug/ml (red curve in **Fig. 2B**), was able to show a complete inhibition curve in the presence of ACE2-IgMu.

**Figure 2.**
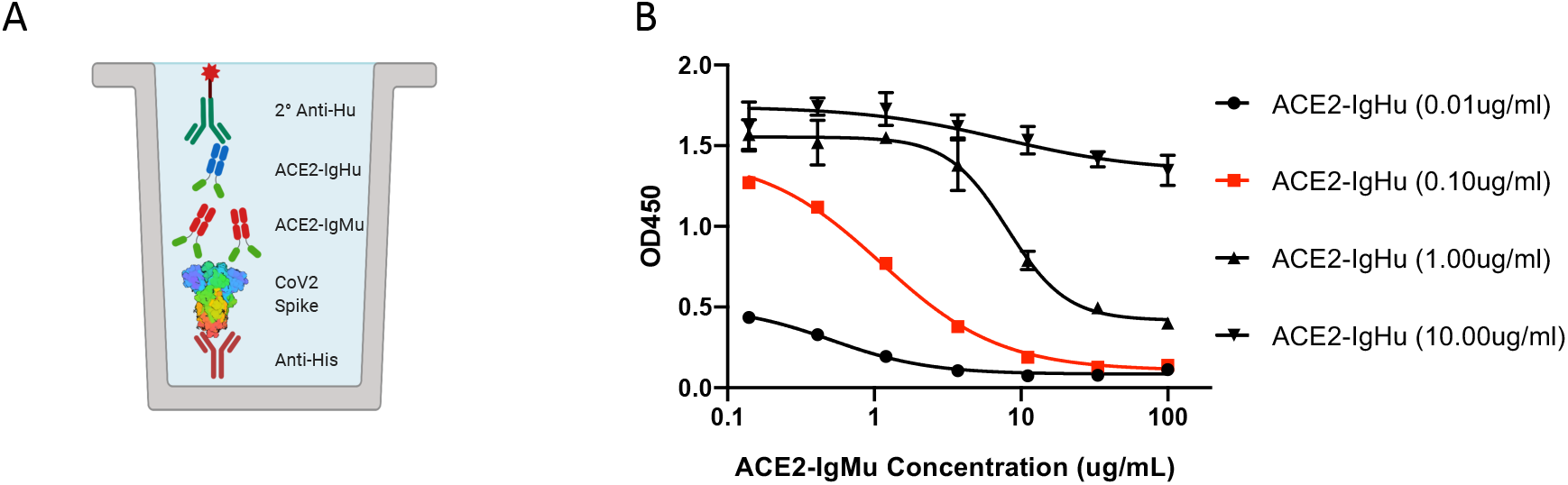
ACE2 receptor competition assay development. **a**) Competition ELISA schematic displaying immobilized anti-His pAb (red) capturing His6X-tagged SARS-CoV-2 spike protein (rainbow). Premixed ACE2-IgHu (green, blue) at a constant concentration with a dilution series of competitors (green, red) is added and anti-human-HRP (green) determines amount of ACE2-IgHu remaining in the presence of competitors through a colorimetric readout **b**) Four constant concentrations of ACE2-IgHu were tested with varying concentrations of the ACE2-IgMu competitor to establish an optimal ACE2-IgHu concentration which displays a full blocking curve (red, 0.10ug/ml) from the competitor dilution series while retaining a wide range in signal.

### Animal IgG and serological competition

The proof-of-concept competition ELISA displayed a full blocking curve, so we sought to utilize this assay for animals immunized with SARS-CoV-2 spike protein. The same design for the competition ELISA was used for this assay, replacing the ACE2-IgMu competitor with antibodies induced by vaccination (**Fig. 3A**). In our previous work, BALB/c mice were immunized with DNA plasmids encoding SARS-CoV-2 spike protein(23). To examine the activity of antibodies in the sera, IgGs from either naïve mice or vaccinated mice 14 days post-immunization were purified using a protein G column. Unlike the ACE2 control which only binds to the receptor binding site (RBS) on the receptor binding domain (RBD) of the spike protein, antibodies from immunized mice can bind to a multitude of epitopes on the spike protein including epitopes on the S1 subunit(which includes the RBD), S2 subunit or S1-S2 interfaces. While antibodies distal to the RBS should have less effect on ACE2 binding, we hypothesized that such distal antibodies may inhibit ACE2 binding directly by sterically obscuring the RBS or indirectly by causing allosteric conformational shifts in the spike protein. To test this, we immobilized either the full spike protein (S1+S2) or S1 alone to examine the levels of detectable blocking antibodies. A mixture of ACE2-IgHu at a constant concentration of 0.1 ug/ml and a dilution series from a vaccinated mouse IgG (IgGm1) or naïve mice IgG (naïve IgGm) was incubated on the plate. An anti-human-HRP conjugated antibody was added to determine the ACE2 binding in the presence of IgGm. As **Fig. 3B** illustrates, there is greater antibody blocking with the full spike protein than with the S1 subunit alone (Fig. S2A).

**Figure 3.**
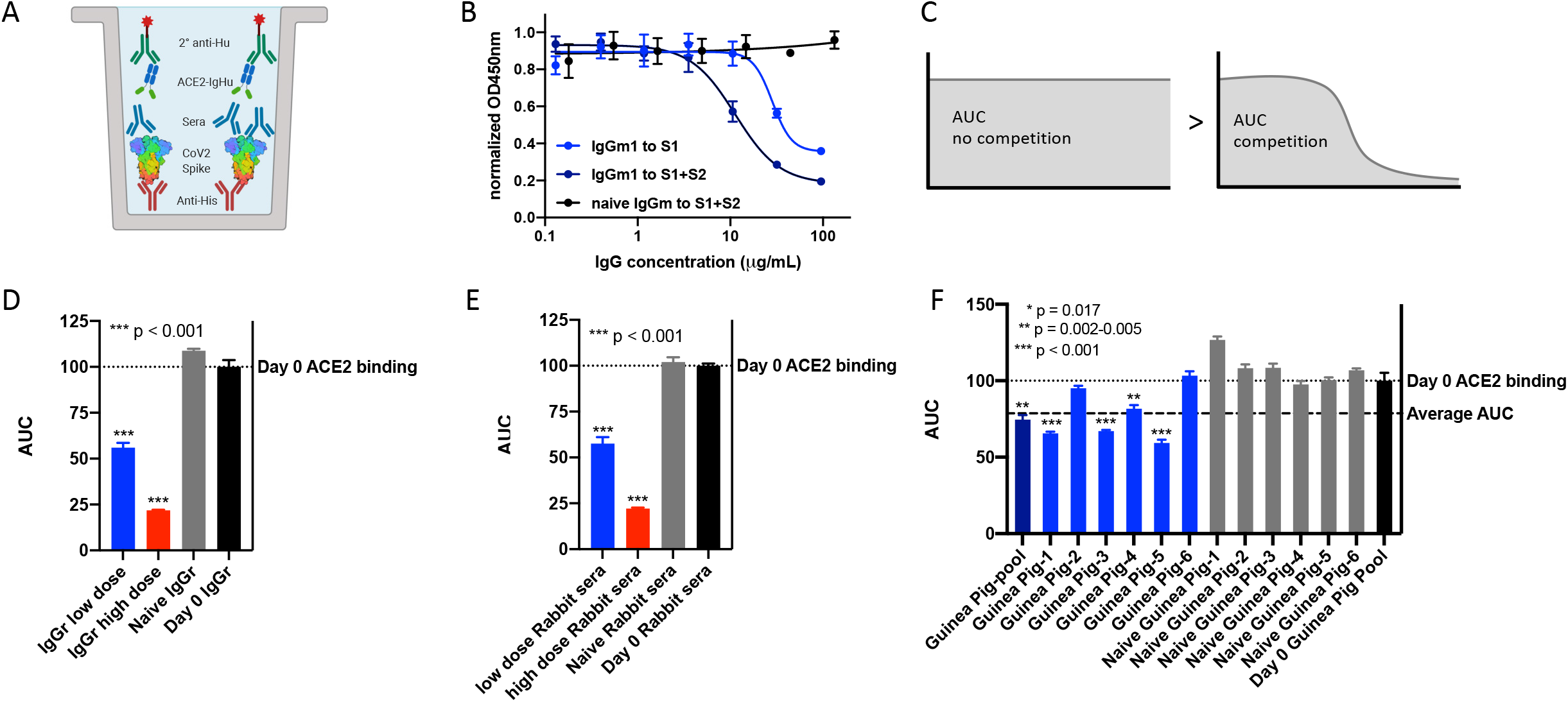
Animal IgG and serological competition. **a**) IgG and serological competition schematic. Anti-his pAb captures CoV-2 spike protein. Immunized sera or IgG from small animals are used as competitors to block ACE2-IgHu receptor binding when premixed. ACE2-IgHu remaining is determined from an anti-human-HRP colorimetric readout **b**) IgGs present in a vaccinated BALB/c mouse block ACE2-IgHu binding with greater effect when full-length SARS-CoV-2 S1-S2 spike protein is immobilized versus the S1 subunit by itself **c**) Area Under the Curve (AUC) schematic displaying the larger area for uninhibited ACE2 binding versus the area from curves showing competition with ACE2 **d**) AUC of IgGs purified from immunized rabbit sera (IgGr low dose, blue; IgGr high dose, red) vs naïve IgGr or Day 0 IgGr. **e**) AUC of sera from immunized rabbits (low dose Rabbit sera, blue; high dose Rabbi sera, red) versus naïve Rabbit sera or Day 0 Rabbit sera. **f**) AUC of sera from immunized guinea pigs at week 2 (dark blue) and individual animals (blue), naïve sera (grey) and pooled Day 0 sera from all animals (black). The pooled immunized curve displayed a comparable AUC to the average AUC from all individual immunized animals.

To show the utility of this assay in samples from larger mammals, we examined receptor blocking of rabbit sera from SARS-CoV-2 immunization studies. We pooled sera from five rabbits two weeks post immunization in 3 groups (low dose of 1mg, high dose of 2mg and naïve) as well as a Day 0 pool from all 15 animals. The IgG from these pools were purified on a protein G column and used as competitors in the competition ELISA assay. To compare the blocking efficacy, the area under the curve (AUC) was calculated for each competition curve. For full, uninhibited ACE2 binding, the AUC will be larger than the AUC for a competitive curve (**Fig. 3C**). As seen with the mouse IgGs, the pooled vaccinated rabbit IgG displayed statistically significant blocking of ACE2 receptor binding compared to the naïve animal pool and the Day 0 pool (**Fig. 3D, Fig S2B**). The high dose rabbits reduced ACE2 signal relative to the low dose group, highlighting the utility of the assay to help discriminate between different vaccine regimens. Up to this point we have analyzed purified IgGs collected from sera, however we wanted to validate the use of this assay on serological samples as well. The same rabbit sera pools were used as competitors in the competition ELISA assay in a dilution series to compare blocking between sera and purified IgG. The rabbit sera displayed statistically significant ACE2 receptor blocking as we saw in the purified IgG assay (**Fig. 3E, Fig. S2C**).

Next, we sought to show that we could assess receptor blocking in a third animal model of guinea pigs which were immunized with a SARS-CoV-2 Spike-based vaccine(23). We compared a pool of sera collected on day 14 post immunization to a Day 0 pool (**Fig. 3F**). The immune pool showed significantly lower AUC signal indicating ACE2 blocking ability of the immune sera. We then sought to compare how animal sera pools represent blocking ability of the individual animals. The AUC was calculated for each normalized curve and plotted with the immunized guinea pig pool AUC. Four of the six guinea pigs immunized against SARS-CoV-2 spike showed statistically significant ACE2 blocking and, importantly, the pooled sera was comparable to the average AUC from all six sera samples (**Fig. 3F, Fig. S2D**). The competition ELISA was used to analyze both the IgGs and the sera from the groups, both groups showed statistically significant blocking of the ACE2 receptor in these assays (**Fig. S2E, Fig. S2F**).

### Primate serological competition

With the competition ELISA assay demonstrated to be capable of measuring molecular blocking of the ACE2 receptor in small animals with sera and IgG, we next altered the assay for use with primate samples. In the small animal studies, the anti-human secondary was used to detect soluble receptor ACE2-IgHu. However, this secondary antibody would cross-react with antibodies from primates. To remedy this, an ACE2-mouse Fc fusion (ACE2-IgMu) was utilized in place of the ACE2-IgHu and an anti-mouse Fc secondary antibody HRP conjugate was used for detection of ACE2 binding in the presence of primate antibody inhibitors (**Fig. 4A**). To confirm the function of the replacement ACE2-IgMu, an initial binding ELISA was performed (**Fig. 4B**). We predicted a similar concentration was needed for optimal competition on the binding curve (**Fig. 4B, red arrow**) and confirmed this by running a competition ELISA assay using ACE2-IgHu as the competitor at varying ACE2-IgMu constant concentrations (**Fig. 4C, Fig. S3A**). NHP sera from five animals immunized with SARS-CoV-2 spike were pooled and Day 0 sera from the same group of five animals were pooled as a negative control. The NHP immune sera displayed appreciable ACE2 blocking compared to the naïve sera (**Fig 4D, Fig. S3B**). Next, we examined if we could detect the presence of receptor blocking antibodies in a human ICU patient with acute infection. We compared nine positive COVID-19 patients with sixteen naïve donor samples taken from three years prior to the pandemic. The sera from the COVID-19 patients was able to block ACE2 receptor binding with statistical significance (**Fig. 4E, Fig. S4A**). To compare healthy and COVID-19 patient samples in an independent assay, we employed a pseudovirus assay we recently developed(23). Sera from the healthy donors could not neutralize the virus, yet sera from COVID-19 patient samples could neutralize the virus (**Fig. 4F, Fig S4B, S4C**). This finding is consistent with the ACE2 blocking data and we show a correlation between pseudovirus neutralization ID50 and AUC for residual ACE2 blocking across all our datasets (**Fig. 5**). Thus, we have demonstrated that the ACE2 competition assay can be employed to measure receptor inhibition levels of human samples.

**Figure 4.**
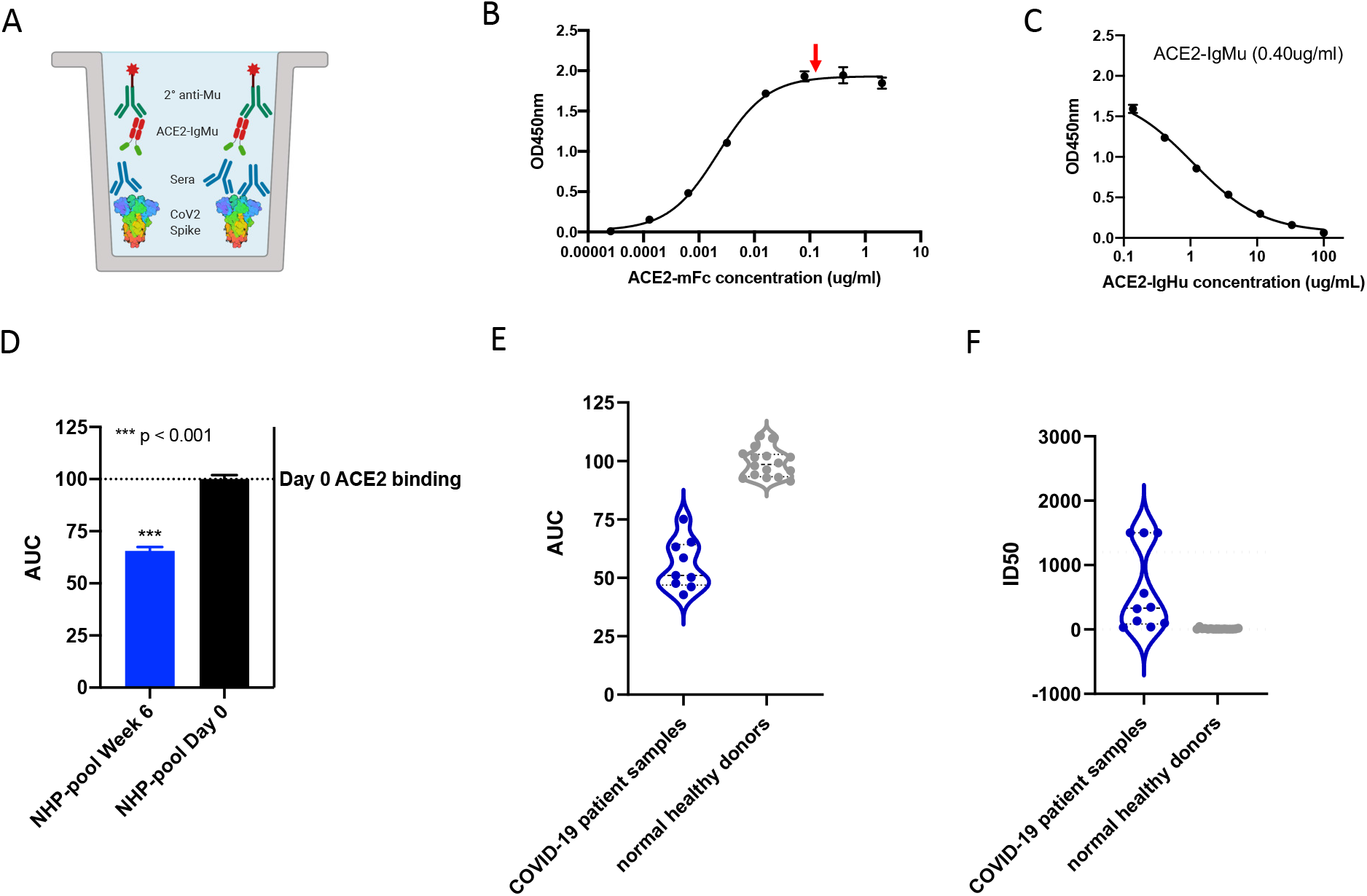
Primate serological competition. **a)** Competition ELISA schematic displaying immobilized His6X-tagged CoV-2 spike protein (rainbow). Preblocking of the spike protein with primate sera (blue) at varying concentrations was added followed by ACE2-IgMu (green, blue) at a constant concentration. Anti-mouse-HRP (green) determines amount of ACE2-IgMu remaining in the presence of competitors through a colorimetric readout **b)** Affinity of ACE2-IgMu for immobilized SARS CoV-2 S1+S2 full-length spike protein assessed by ELISA. Optimal concentration of ACE2-IgMu for competition assays (red arrow, 0.4ug/ml) requires high signal without excess receptor present**. c)** Optimal ACE2-IgMu concentration which displays a full blocking curve (0.40ug/ml) from the competitor dilution series (ACE2-IgHu) while retaining a wide range in signal **d)** NHP sera pooled from five vaccinated animals was used as competitors in the primate competition assay. The AUC from vaccinated NHP sera (blue) versus Day 0 NHP sera (black). **e)** Human sera from nine SARS-CoV-2 positive COVID-19 patients was tested in the primate competition assay and compared to sixteen naïve human sera collected pre-pandemic. The AUC of the COVID-19 patient serum (purple) is significantly decreased compared to the pre-pandemic human serum (grey). **f)** Human sera was analyzed by a pseudovirus neutralization assay. The samples and the coloring are the same as in e).

**Figure 5.**
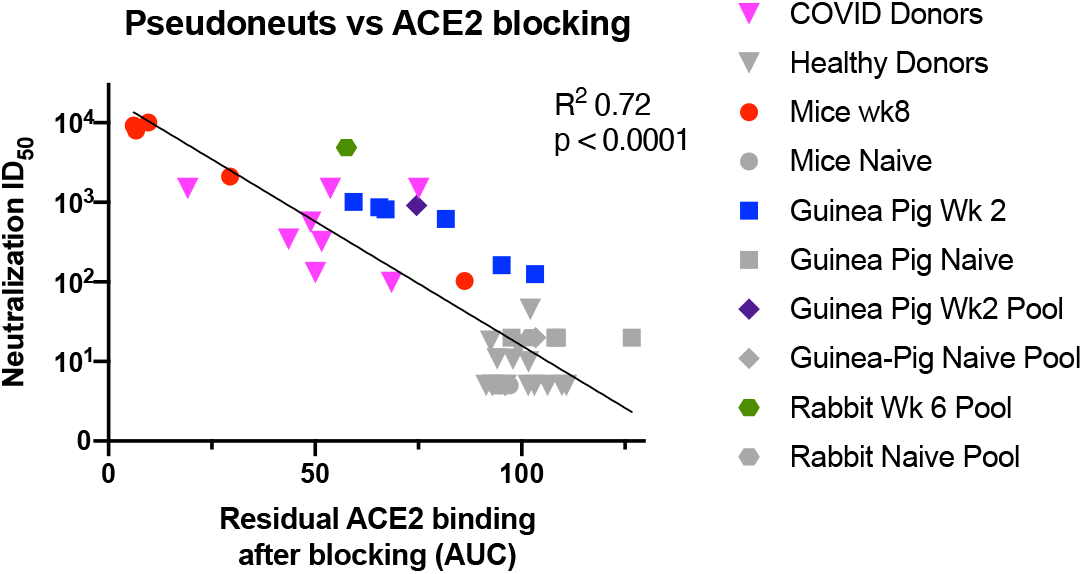
ACE2 receptor blocking correlates with psuedovirus neuralization. A symbol represents each of the individual datapoints where we had a paired AUC blocking and pseudovirus ID50 values. The human samples are in triangles, the mice in circles, individual Guniea pigs in squares, Guniea pig pools in diamonds and rabbit pools in hexagons. SARS-CoV-2 spike experienced samples are shown in color. Naïve samples and healthy donors are shown in grey. Least-squares fit line is shown with p-value and R squared from Prism.

### SPR-based assay for ACE2 receptor blocking

To quantitate blocking of the Spike-ACE2 bimolecular interaction in a second, independent experiment, we developed a sensitive surface plasmon resonance (SPR) assay. SPR is a widely used platform that does not require secondary antibodies and therefore we could use a single assay format for small animals, NHPs and humans. In our assay, a CAP sensor chip is used to capture single stranded DNA coupled to streptavidin. We are then able to capture biotinylated spike protein to the surface of the sensor chip (**Fig. 6A**). In SPR, changes in refractive index occurs when molecules interact with the sensor surface or proteins attach to the sensor surface and these changes are reported as resonance units (RUs). Next, anti-spike samples in a dilution series can be injected. After a short time, ACE2 receptor at a constant concentration can be injected to measure residual receptor binding. The change in RUs after the second injection are a measure of ACE2 blocking.

**Figure 6.**
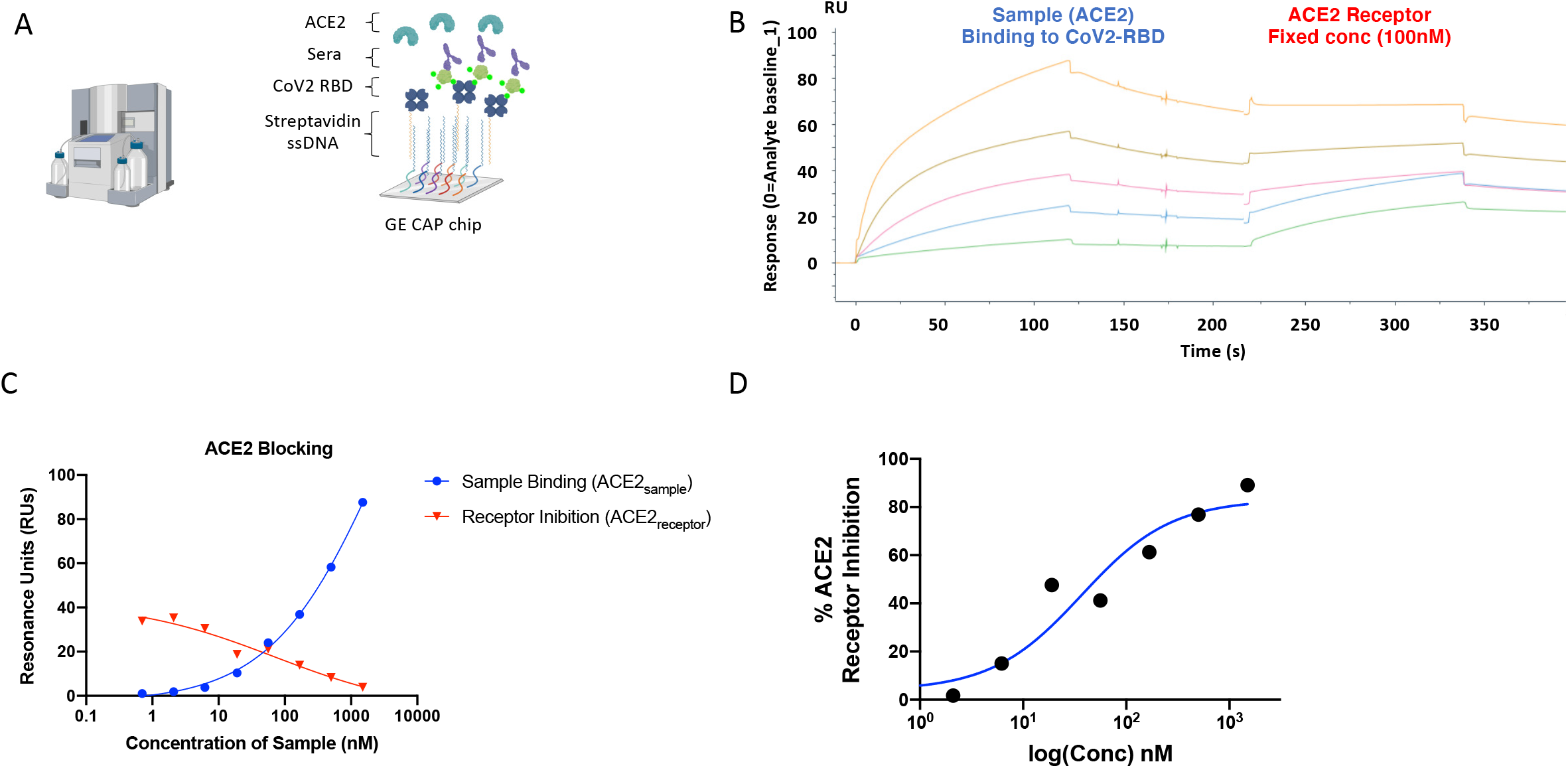
ACE2 receptor competition assay development. **a)** Overview of SPR experiment depicting SARS-CoV-2 capture by streptavidin-biotin interaction, sera injected as analyte and ACE2 injected as second analyte **b)** Sensorgram for ACE2 blocking SPR assay ACE2-IgHu injected as sample as indicated and ACE-IgHu injected as receptor as indicated. Sample responses were referenced to blank injections. **c)** Response in RUs measured at the end of sample injection (blue) and receptor injection (red) at each concentration of sample. **d)** ACE2 inhibition curve derived from RUs at each concentration

To demonstrate the feasibility of this assay, we used ACE2-IgHu as both the sample (ACE2_sample_) and the receptor (ACE2_receptor_). The sensorgram for this experiment shows ACE2_sample_ injections at various concentrations binding to SARS-CoV-2 RBD between 0 and 125 seconds (**Fig. 6B**). At 225 seconds we inject ACE2_receptor_ at 100nM. We observe ACE2_receptor_ binding to SARS-CoV-2 RBD at the lower ACE2_sample_ concentrations, but not at the highest ACE2_sample_ concentration. The binding signals of ACE2_sample_ and ACE2_receptor_ intersect close to 100nM, suggesting the assay is working as expected (**Fig 6C**). A measure of % ACE2 receptor inhibition can be calculated to the dose dependence response of the sample (**Fig. 6D**). While this experiment was done with samples containing a human Fc, this assay can be employed to measure inhibition of the spike:receptor interaction with any sample regardless of species.

## Discussion

Diagnostic methods that fast and accurately detect functional immunity in a SARS-CoV-2 patient or vaccinee are of critical importance during the global pandemic of 2020. Serological tests are a central component of our SARS-CoV-2 diagnostic toolbox. Serological tests are often configured to measure the presence of antibodies that recognize SARS-CoV-2 proteins. This is not a sufficient measurement to determine if a donor’s blood should be used to treat seriously ill patients or to determine if the donor could be re-infected. Instead, a measurement of functional antibodies is required. Further, systematic studies of functional antibody responses and persistence in patients presenting asymptomatic, mild and severe cases will be critical to understand SARS-CoV-2 immunity. In this study, we developed two new assays which can aid in understanding the specificity and functionality of antibodies in serum. First, we showed that an ELISA-based assay could be employed to measure receptor blocking antibodies in animals vaccinated with SARS-CoV-2 spike protein. We could measure purified IgG or directly from sera, as pools across groups of animals or individual animals. The assay is robust as exemplified by the ease of adaption to use for five different species. Importantly, the values of ACE2 blocking measured in our assay correlate with pseudovirus neutralization titers, as seen in Figure 5. Second, we developed an SPR assay for detecting inhibition of the ACE2 receptor binding to spike protein.

Our assays focus on blocking receptor interactions as a surrogate for virus neutralization. In this study we studied ACE2, because ACE2 appears to be the major receptor for SARS-CoV-2. The assay could be also be used to study antibodies targeting other coronaviruses with similar receptors such as SARS-CoV-1 or NL63. The Basigin receptor, as known as CD147, has been demonstrated to play a role in SARS-CoV-1 infection(24) and is now hypothesized to function in SARS-CoV-2 invasion of host cells(25). By engineering the ectodomain of CD147 with human and murine Fc tags, these assays could easily be adapted to study antibody inhibition of CD147. In addition, antibodies against other coronaviruses, such as MERS, which utilize the DPP4 receptor could be studied in our assay format using a similar adaptation.

Neutralizing antibodies can target viral surface proteins to inhibit virus function. Recently, an antibody (B38) was discovered that can neutralize SARS-CoV-2 and has been structurally defined to bind to the same site as ACE2(26). In addition, three other antibodies (H4, B5, H2) all showed neutralizing activity, but B5 and H2 did not compete ACE2 completely, suggesting they bind an alternative neutralizing epitope on the RBD. In another recent study, a nAb, C12.1, binds the RBD spike protein, potently neutralizes and protects Syrian hamsters compared to controls in a passive transfer experiment(27). Recently, a human monoclonal antibody isolated from transgenic mice (47D11) has been shown to neutralize both SARS-CoV-1 and SARS-CoV-2(28). Probing serological responses for competition and blocking of these protective epitopes could also be easily accomplished through the ELISA and SPR assays reported here.

The Spike-ACE2 interaction is also being considered as an important therapeutic target. Indeed, there have been SARS-CoV-1 Spike-ACE2 inhibitors developed previously(29). To examine the functionality of these small molecules beyond direct binding to the spike protein, assays such as the one developed here are needed. The SPR instrument is often used for drug discovery and the SPR assay could be easily adapted to examine blocking capabilities of candidate drugs. The ELISA assay does not depend on the molecular identity of the competitor, so small molecule or peptide inhibitors could be directly assessed in this assay.

Our study presents a new set of assays for assessing ability of antibody samples to inhibit SARS-CoV-2 Spike interaction with its receptor. As with most assays, the limit of detection can be an issue. In some of our samples, we saw robust blocking and in others there was minimal blocking. This could be a property of the samples themselves or a limit in the ability to detect ACE2 inhibition in our assays. In addition, discovering functional monoclonal antibodies can be important to understand SARS-CoV-2 immunity and serological assays of functional antibodies is just a first step. Indeed, a recent report observed that only a small portion of monoclonal antibodies sorted from convalescent donors are neutralizing(27). Therefore, it is important to continue to develop and publish assays of varying formats to detect functional antibody responses in SARS-CoV-2 spike exposed people.

## Conclusions

In a fast-moving global pandemic, we must quickly bring to bear all our immunological knowledge to create new tools. Receptor-blocking assays can detect functional antibodies in serum samples. Functional antibody assays can be employed widely to help study SARS-CoV-2 infection, create ‘immune passports’ to enable safe exit from quarantine and for assessing efficacy of vaccines.

## Methods

### DNA design and plasmid synthesis

Protein sequences for human Angiotensin-converting enzyme 2 (huACE2) were obtained from UniProt (Q9BYF1). DNA encoding the IgG1 human Fc sequence was added to the C-terminus of the huACE2 and cloned by Twist Bioscience into a modified mammalian pVax-1 expression plasmid to generate the recombinant plasmid, huACE2-IgHu-pVax.

### Recombinant protein expression and purification

Expi293F cells were transfected with plasmid expressing huACE2-IgHu-pVax using PEI/Opti-MEM. Cell supernatants were harvested 6 days post-transfection by centrifuging (4000xg, 25mins) and filtering (0.2um Nalgene Rapid-flow filter). Supernatants were first purified with affinity chromatography using AKTA pure 25 system and HiTrap MabSelect SuRe protein A column (GE healthcare, Cat# 11-0034-94) for huACE2-IgHu-pVax. The eluate fractions from the affinity purification were pooled, concentrated and dialyzed into 1x PBS buffer before being further purified by size exclusion chromatography (SEC) using Superdex 200 10/300 GL column (GE healthcare). Identified SEC eluate fractions were pooled and concentrated to 1mg/mL. The molecular weight and homogeneity of the purified sample were confirmed through size exclusion chromatography multi-angle light scattering (SEC MALS) by running the sample in PBS over Superose 6 increase column followed by DAWN HELEOS II and Optilab T-rex detectors. The data collected was analyzed using the protein conjugated analysis from ASTRA software (Wyatt technology).

#### Biophysical characterization of ACE2-IgHu

SPR experiments were performed using a protein A capture chip on a Biacore 8k. The running buffer was HBS-EP with 1mg/ml BSA. The SARS-CoV-2 RBD was used as an analyte at concentrations of 100nM, 10nM and 1nM. The experiment had an association phase for 120sec and the dissociation phase for 600sec. The data was fit by a 1:1 Langmuir model.

##### SARS-CoV2 Spike binding huACE2-IgHu/ huACE2-IgMu ELISA

96-well half area plates (Corning) were coated at room temperature for 8 hours with 1ug/mL PolyRab anti-His antibody (ThermoFisher, PA1-983B), followed by overnight blocking with blocking buffer containing 1x PBS, 5% skim milk, 1% FBS, and 0.2% Tween-20. The plates were then incubated with 10ug/mL of SARS-CoV-2, S1+S2 ECD (Sino Biological, 40589-V08B1) at room temperature for 1-2 hours, followed by addition of either huACE2-IgHu serially diluted 5-fold (starting concentration, 2ug/mL) or huACE2-IgMu (Sino Biological, 10108-H05H) serially diluted 5-fold (starting concentration, 50ug/mL). Serial dilutions were performed using PBS with 1% FBS and 0.2% Tween and incubated at RT for 1 hour. The plates with huACE2-IgHu were then incubated at room temperature for 1 hour with goat anti-human IgG-Fc fragment cross adsorbed Ab (Bethyl Laboratories, A80-340P) at 1:10,000 dilution. Similarly, the plates with huACE2-IgMu were incubated with goat anti-mouse IgG H+L HRP (Bethyl Laboratories, A90-116P) at 1:20,000 dilution. Next, TMB substrate (ThermoFisher) was added to both plates and then quenched with 1M H_2_SO_4_. Absorbance at 450nm and 570nm were recorded with a BioTek plate reader. Four washes were performed between every incubation using PBS with 0.05% Tween.

##### Competition ELISA- Control (huACE2-IgMu Vs huACE2-IgHu)

96-well half area plates (Corning) were coated at room temperature for 8 hours with 1ug/mL PolyRab anti-His antibody (ThermoFisher, PA1-983B), followed by overnight blocking with blocking buffer containing 1x PBS, 5% skim milk, 1% FBS, and 0.2% Tween-20. The plates were then incubated with 10ug/mL of SARS-CoV-2, S1+S2 ECD (Sino Biological, 40589-V08B1) at room temperature for 1-2 hours. huACE2-IgMu (Sino Biological, 10108-H05H) was serially diluted 3-fold (starting concentration,100ug/mL) with PBS with 1% FBS and 0.2% Tween and pre-mixed with huACE2-IgHu at constant concentrations (ranging from 0.01-10ug/mL) The pre-mixture was then added to the plate and incubated at RT for 1 hour. The plates were further incubated at room temperature for 1 hour with goat anti-human IgG-Fc fragment cross adsorbed Ab (Bethyl Laboratories, A80-340P) at 1:10,000 dilution, followed by addition of TMB substrate (ThermoFisher) and then quenched with 1M H_2_SO_4_. Absorbance at 450nm and 570nm were recorded with a BioTek plate reader. Four washes were performed between every incubation using PBS with 0.05% Tween.

##### Competition ELISA- Mouse

IgG was purified from sera taken from mice immunized with SARS-CoV-2 Spike protein (INO-4800) (23) using Nab protein G spin kit (ThermoFisher, 89949). 96-well half area plates (Corning) were coated at room temperature for 8 hours with 1ug/mL PolyRab anti-His antibody (ThermoFisher, PA1-983B), followed by overnight blocking with blocking buffer containing 1x PBS, 5% skim milk, 1% FBS, and 0.2% Tween-20. The plates were then incubated with 10ug/mL of SARS-CoV-2, S1+S2 ECD (Sino Biological, 40589-V08B1) at room temperature for 1-2 hours. Purified IgG was serially diluted 3-fold (starting concentration,100ug/mL) with PBS with 1% FBS and 0.2% Tween and pre-mixed with huACE2-IgHu at constant concentration of 0.1ug/mL. The pre-mixture was then added to the plate and incubated at RT for 1 hour. The plates were further incubated at room temperature for 1 hour with goat anti-human IgG-Fc fragment cross adsorbed Ab (Bethyl Laboratories, A80-340P) at 1:10,000 dilution, followed by addition of TMB substrate (ThermoFisher) and then quenched with 1M H_2_SO_4_. Absorbance at 450nm and 570nm were recorded with a BioTek plate reader. Four washes were performed between every incubation using PBS with 0.05% Tween.

##### Competition ELISA- Guinea pig

Sera from guinea pigs immunized with INO-4800(23) was collected and pooled together prior to immunization (Day 0, n=5) and two weeks post immunization. Sera from naïve guinea pigs (n=5) given a saline control was also pooled. IgG was purified from these sera pools using Nab protein A/G spin kit (Cat# Thermo, 89950). 96-well half area plates (Corning) were coated at room temperature for 8 hours with 1ug/mL PolyRab anti-His antibody (ThermoFisher, PA1-983B), followed by overnight blocking with blocking buffer containing 1x PBS, 5% skim milk, 1% FBS, and 0.2% Tween-20. The plates were then incubated with 10ug/mL of SARS-CoV-2, S1+S2 ECD (Sino Biological, 40589-V08B1) at room temperature for 1-2 hours. Either purified sera or IgG (Day0, Day14 or naïve) was serially diluted 3-fold with PBS with 1% FBS and 0.2% Tween and pre-mixed with huACE2-IgHu at a constant concentration of 0.1ug/mL. The pre-mixture was then added to the plate and incubated at RT for 1 hour. The plates were further incubated at room temperature for 1 hour with goat anti-human IgG-Fc fragment cross adsorbed Ab (Bethyl Laboratories, A80-340P) at 1:10,000 dilution, followed by addition of TMB substrate (ThermoFisher) and then quenched with 1M H_2_SO_4_. Absorbance at 450nm and 570nm were recorded with a BioTek plate reader. Four washes were performed between every incubation using PBS with 0.05% Tween.

##### Competition ELISA- Rabbit

Rabbits were immunized with 1mg or 2mg of ION-4800 at Day 0 and Week 4. Sera from immunized rabbits was collected and pooled together prior to immunization (Day 0, n=5) and 2 weeks post 2nd immunization (wk 6, n=5), (Patel A. et al., in preparation). Sera from naïve rabbits (n=5) given a saline control was also pooled. IgG was purified from these sera pools using Nab protein A/G spin kit (Thermo, 89950). 96-well half area plates (Corning) were coated at room temperature for 8 hours with 1ug/mL PolyRab anti-His antibody (ThermoFisher, PA1-983B), followed by overnight blocking with blocking buffer containing 1x PBS, 5% skim milk, 1% FBS, and 0.2% Tween-20. The plates were then incubated with 10ug/mL of SARS-CoV-2, S1+S2 ECD (Sino Biological, 40589-V08B1) at room temperature for 1-2 hours. Either sera pool or purified IgG (IgGr) from Day0, low dose, high dose or naïve animals was serially diluted 3-fold with PBS with 1% FBS and 0.2% Tween and pre-mixed with huACE2-IgHu at constant concentration of 0.1ug/mL. The pre-mixture was then added to the plate and incubated at RT for 1 hour. The plates were further incubated at room temperature for 1 hour with goat anti-human IgG-Fc fragment cross adsorbed Ab (Bethyl Laboratories, A80-340P) at 1:10,000 dilution, followed by addition of TMB substrates (ThermoFisher) and then quenched with 1M H_2_SO_4_. Absorbance at 450nm and 570nm were recorded with a BioTek plate reader. Four washes were performed between every incubation using PBS with 0.05% Tween.

##### Competition ELISA- Primates

Rhesus macaques were immunized with 1mg of INO-4800 at Day 0 and Week 4. Sera from immunized NHPs was collected and pooled together prior to immunization (Day 0, n=5) and 2 weeks post 2nd immunization (Wk 6, n=5), (Patel A. et al., in preparation). Sera from SARS-CoV-2 positive patients treated at the Hospital of the University of Pennsylvania was also collected and compared to sera from heathly human donors purchased from BioChemed in December of 2016, well before the SARS-CoV-2 pandemic. 96-well half area plates (Corning) were coated at 4°C overnight with 1ug/mL of SARS-CoV-2, S1+S2 ECD (Sino Biological, 40589-V08B1). Plates were washed 4x with 1x PBS, 0.05% Tween-20 using a plate washer, and then blocked with 1x PBS, 5% skim milk, 0.1% Tween-20 at room temperature for 1 hour. After washing, sera was serially diluted 3-fold in 1x PBS, 5% skim milk, 0.1% Tween-20 and incubated on the plate at room temperature for 1 hour. Plates were washed, then incubated with a constant concentration of 0.4 ug/mL huACE2-IgMu diluted in 1x PBS, 5% skim milk, 0.1% Tween-20 at room temperature for 1 hour. After washing, plates were further incubated at room temperature for 1 hour with goat anti-mouse IgG H+L HRP (Bethyl Laboratories, A90-116P) at 1:20,000 dilution in 1x PBS, 5% skim milk, 0.1% Tween-20. Plates were then washed, followed by addition of TMB substrate (ThermoFisher) and then quenched with 1M H_2_SO_4_. Absorbance at 450nm and 570nm were recorded with a BioTek plate reader. Four washes were performed between every incubation using PBS with 0.05% Tween.

### SPR Assay

Surface Plasmon Resonance experiments were conducted on a Biacore 8K instrument. The running and dilution buffers were HBS-EP, 1 mg/ml BSA, 0.05% Tween. Biotin CAPture reagent was injected over the Series S CAP Sensor chip (Cytiva Life Sciences 28920234) at 2 ul/min for 300s. Followed by capture of SARS-CoV-2 receptor binding domain which was biotinylated using the lightning-link type-A biotinylation kit(Expedeon/Abcam, 370-0005) for 180s at 10ul/min. Next, the sample (ACE2-IgHu) at 8 concentrations of 1500nM to 0.7nM. The association was for 120s second at a flow rate of 30 ul/min and followed by a short dissociation time of 15s. A constant ACE2-IgHu was injected at 100 nM in each flow cell for 120s followed by 60s dissociation at a flow rate of 30ul/min. The difference in response units before injection and after dissociation of the sample application and the constant receptor was calculated for each curve. The regeneration solution was made using 3-parts 8M guanidine hydrochloride and 1-part 1M sodium hydroxide.

### Statistics

Statistical analyses were performed using GraphPad Prism 8 software. The data were considered significant if p <0.05. The lines in all graphs represent the mean value and error bars represent the standard deviation. The AUC values were normalized as a percentage of Day 0 AUC. No samples were excluded from the analysis. Samples were not blinded before performing each experiment.

### SARS-CoV-2 Pseudovirus assay

The SARS-CoV-2 pseudovirus was produced by co-transfection of HEK293T cells with 1:1 ratio of DNA plasmid encoding SARS-CoV-2 S protein (GenScript) and backbone plasmid pNL4-3.Luc.R-E-(NIH AIDS Reagent) using GeneJammer (Agilent) in 10% FBS/DMEM enriched with 1% Pennicilin/Streptomycin (D10 medium). The supernatant containing pseudovirus was harvested 48 hours post-transfection and enriched with FBS to 12% total volume, sterifiltered and stored at −80C. The pseudovirus was tittered using a stable ACE2-293T cell line that had previously been generated(23). For neutralization assay, serially diluted samples were incubated with pseudovirus at room temperature for 90 minutes and added to 10,000 ACE2-293T cells in 200uL TPCK media (DMEM supplemented with 1%BSA, 25mM HEPES, 1ug/ml of TPCK and 1X Penicillin-Streptomycin) in 96-well tissue culture plates, and incubated at 37C and 5% CO_2_ for 72 hours. The cells were subsequently harvested and lysed with BriteLite reagent (PerkinElmer), and luminescence from the plates were recorded with a BioTek plate reader.

## Availability of data and materials

The data are available from the corresponding author upon request.

## Abbreviations

## Acknowledgements

The authors would like to Matthew Sullivan for providing feedback on the manuscript.

## Contributions

S.W., N.C. and D.W.K.: Developed the receptor-blocking assays. Designed and characterized ACE2-IgHu protein and the SARS-CoV-2 RBD protein. Planned and executed the experiments and analyzed the data. E.L.R. and K.Y.K.: optimized, planned and executed the experiments involving the human samples. E.L.R., E.G., A.P., P.T.: produced reagents. M. P. and Z.X. created and executed the pseudovirus neutralization assay. S.W., N.C. and D.W.K. wrote the paper. E.L.R., E.G., M.P., Z.X., A.P., D.B.W. helped write the paper.

## Declarations

## Ethics approval and consent to participate

N/A - This study did not require ethics approval and consent.

## Consent for publication

N/A – no identifiable information was included.

## Additional information

## Supplemental Figs

**Supplemental Figure 1. ACE2 receptor expression and assay development.** a) ACE2-IgHu binding comparison for immobilized S1 spike protein from varying batches of expression and purification as well as freeze-thaw assessment via ELISA show negligible differences.

**Supplemental Figure 2. Animal IgG and serological competition. a**) AUC is significantly decreased in the presence of vaccinated mouse IgG competitors; however a greater decrease is observed when full-length CoV-2 spike protein was immobilized versus naïve mouse IgG samples. **b**) ELISA competition curves for vaccinated rabbit IgG (IgGr low dose, blue; IgGr high dose, red) or sera (sera low dose, blue; sera high dose, red) versus naïve rabbit IgG or **c)** sera samples (grey) and pooled Day 0 rabbit IgG or sera samples (black). **d**) ELISA competition curves for Week 2 vaccinated guinea pig sera (pool, dark blue; individual animals, blue) versus naïve guinea pig sera samples (grey) and pooled prevaccinated guinea pig sera samples (black). **e**) AUC for vaccinated guinea pig IgG pool (blue) versus naïve guinea pig IgG pool (grey) and pooled prevaccinated guinea pig IgG samples (black) **f)** ELISA competition curves with same coloring as in e.

**Supplemental Figure 3. Primate serological competition. a**) Four constant concentrations of ACE2-IgMu were tested with varying concentrations of the ACE2-IgHu competitor to establish an optimal ACE2-IgMu concentration which displays a full blocking curve (red, 1.0ug/ml) from the competitor dilution series while retaining a wide range in signal. **b**) ELISA competition curves for vaccinated NHP sera (blue) versus pooled Day 0 NHP sera (black).

**Supplemental Figure 4. a**) Human sera from nine SARS-CoV-2 positive COVID-19 patients was tested in the primate competition assay and compared to sixteen naïve human sera collected pre-pandemic. The AUC of the COVID-19 patient serum (purple) is significantly decreased compared to the pre-pandemic human serum (grey) and normalized to a buffer control. **b**) pseudovirus neutralization assay for the nine SARS-CoV-2 positive COVID-19 patients displayed neutralization for all samples to varying degrees **c**) pseudovirus neutralization assay for the sixteen naïve human sera collected pre-pandemic displayed little to no neutralization for all samples.

